# Identifying potential dopamine receptor D2 antagonist from Rauwolfia serpentina – An in-silico virtual screening approach

**DOI:** 10.1101/2024.10.10.617621

**Authors:** Souvik Chakraborty, Sushmita Bhowmick, Tarasankar Maiti

## Abstract

**Background:** To combat the alarming rise of neuropsychiatric diseases like Schizophrenia, new drugs with fewer side effects have to be identified and characterized for the symptomatic treatment of Scz. Positive symptoms associated with Scz are hallucinations and delusions mainly due to the overactivation of the D2 dopaminergic receptors in the cortical pathways. D2 receptor antagonists identified from *Rauwolfia serpentina* are employed for controlling positive symptoms.

**Materials and methods:** The IMPPAT database was used to identify phytoconstituents from the root of *Rauwolfia serpentina* that can be used for antagonizing D2 dopaminergic receptor. 39 compounds were identified and SDF files were obtained from the PubChem database molecular docking with the modelled dopamine D2 receptor was used as a virtual screening tool followed by ADMET screening of the potential lead compound.

**Results:** After docking analysis, 11 compounds having binding affinity ≥ -9 kcal/mol were shortlisted for ADMET analysis. After analysis, 4 compounds (1,2-dihydrovomilenine, Corynanthine, Rauwolscine, and Yohimbine) were identified as potential leads. Among these 4 compounds, 1,2-dihydrovomelenine has been identified as the lead compound.

**Conclusion:** The present study identified a potential new therapeutic compound that can be used for controlling the positive symptoms of Scz patients in vitro as well as in vivo.

## Introduction

Mental disorders like Schizophrenia (Scz) are associated with dopaminergic D2 receptor overactivity and are positively associated with psychotic disorders such as delusions and hallucinations. About 9.8% of all the mental disorder patients in India are suffering from Scz^1^. The dopaminergic hypothesis of Scz states that the positive symptoms of the disorder are associated with the overactivation of D2 dopaminergic receptors^2^. Overexpression of D2 dopaminergic receptors has been identified in brain imaging of Scz patients. Upregulation of D2 receptors in subcortical pathways and nucleus accumbens in Scz patients is associated with the positive symptoms of the disease and in an imaging study it was identified that there is a 5.8% upregulation of D2 receptors in those patients with no history of anti-psychotic treatment^3,4^. Employing a D2 receptor antagonist for treating the positive symptoms of Scz is a well-tested procedure and many drugs such as haloperidol have been designed for this purpose but the major side effects of these drugs have been the main motivation to develop drugs with fewer side effects and good efficacy^5,6,7^.

In-silico drug designing techniques have been used for the identification of novel drugs for many diseases including psychotic disorders. The computational approach helps in finding the most probable and best candidate drug for testing it in-vitro and in-vivo models and this reduces cost as well as time spent on thousands of possible drug candidates.

Phytoconstituents from plants are a useful source of antipsychotic compounds and have been used for treating patients with psychosis^8^.

*Rauwolfia serpentina* (commonly known as Sarpagandha) is a dicot plant found in sub-Himalayan regions and is rich in indole-based alkaloids. *R. serpentina* has been used for treating high blood pressure probably due to the presence of reserpine, a well-known anti-hypertensive drug^9^. In India, for many years, reserpine obtained from the *R. serpentina* was used for psychotic disorders, fever, and in cases of snakebites^10^. This study aims to identify and repurpose antagonists for the D2 receptor from phytoconstituents from the roots of *R. serpentina* following the Indian Medicinal Plants, Phytochemistry, and Therapeutics (IMPPAT) database^11^. A study showed that *R. serpentina* extracts have antipsychotic properties^12^. Virtual screening of phytoconstituents of *R. serpentina* led to the identification of 11 compounds and after ADMET analysis 4 compounds were identified that crossed every filter and they are, 1,2-dihydrovomilenine (-9.3 kcal/mol), Corynanthine (-9 kcal/mol), Rauwolscine (-9.6 kcal/mol) and Yohimbine (-9.1 kcal/mol). Though the above four molecules have passed all the filters used in this in-silico study, known side effects of Yohimbine, Rauwolscine, and Corynanthine have been established by in-vitro methods^13,14^. Therefore the present study was designed to investigate the presence of a potential D2 receptor antagonist from the phytoconstituents of *R. serpentina* and the identified phytoconstituents according to our study is 1,2-dihydrovomelinine.

## Material and methods

### Modeling of protein of interest

The structure of the human D2 dopaminergic receptor was created using MODELLER (version 10.4) which uses a homology-based approach for designing proteins from templates provided^15^. For this study, the FASTA file for the D2 receptor was obtained from the UniProt database and a BLAST (Basic Local Alignment Search Tool) search was performed where in the database section Protein Data Bank (PDB) and in the algorithm section, blastp (protein-protein blast was chosen for the search. The three pdb files with the PDBID 7JVR, 7DFP, and 6CM4 that were chosen for the template sequence for homology-based protein modelling were chosen based on their query cover, E value, and percentage identity. For homology modelling, those templates should be chosen that have low E values and high percentage identity with the protein sequence that has been used in the pblast search. For these three PDB structures 7JVR, 7DFP, and 6CM4, the percentage query cover percentage were 100%, 92%, and 92% the respective E values for the three PDBs are 0.0 and the percentage identity for these three structures are 100%, 68.70, and 66.97 respectively. Discrete optimized protein energy (DOPE) is a combination of statistical scoring functions that that help in determining the energy of a given protein structure based on statistical distribution observed in known protein structure. Protein models with lower DOPE score is considered energetically more suitable model because of low steric clashes and correct torsion angles and bond angles. Based on the lowest DOPE score The best model with the lowest DOPE score was chosen and by using the Ramachandran plot analysis the 3D model was validated using PROCHECK which is a program to check the quality of a protein structure^16^. This program is integrated into a webserver called SAVES (version 6.0) (available at *https://saves.mbi.ucla.edu/*).

### Selection of the compounds

The Indian Medicinal Plants Phytochemistry and Therapeutics (IMPPAT) database (available at *https://cb.imsc.res.in/imppat/home*) was used to identify the root extracts of *R. serpentina* was considered to be useful for Scz treatment^11^. A total of 39 compounds were identified using the IMPPAT database from the root extracts of the *R. serpentina* plant. The SDF files for the identified compounds were downloaded from the PubChem database^17^.

### Virtual screening of the compounds

After uploading the protein structure and ligand in SDF format, energy minimization of all the ligands was performed using an applied universal force field using Open Babel. Both protein and ligands were converted to AutoDock ligand format (PDBQT) and then molecular docking was performed. For screening all the 39 compounds, blind docking was executed using the open access tool PyRx (version 0.8) that uses the AutoDock Vina plugin, and for this study, a grid box of Center X: 23.3397, Y:11.5667, Z: -13.2207 and Dimensions (in Angstrom) of X:100.1366, Y:94.1255, Z:119.9587 that covers the whole protein structure^18^. The output files generated after the docking was complete contain information on different poses of the ligand with their corresponding binding affinity. The docked complexes were visualized using the BIOVIA Discovery studio visualizer to understand the nature of interactions between the ligands and the protein^19^. The binding pose describes the orientation of the ligands in the binding site of the protein and is often represented by the root mean square deviation between the docked pose of the ligand and its experimentally determined pose.

### Bioavailability Radar and testing in-silico toxicity of compounds

Drug likeliness of the identified and virtually screened compounds was studied using the SwissADME webserver (*http://www.swissadme.ch/*)^20^. Bioavailability radar shows six basic biophysical properties of the identified compounds such as size, solubility, polarity, lipophilicity, flexibility, and saturation. Lipinski’s rule of five together with the compound's ability to cross blood blood-brain barrier and also its ability to be absorbed by the GI tract was used as an effective screening methodology for the organic compounds. To check whether these screened compounds have any toxic, tumourigenic, or mutagenic effects, OSIRIS an open-source software was used^21^.

### Molecular dynamic simulations

Normal mode analysis was performed for the 1,2-dihydrovomilenine docked D2 receptor complex. The iMODS webserver was used for this purpose to study the collective functional motions of the biological molecules using normal mode analysis^22^. Deformability and beta factor calculations are one of the few parameters that the iMODS predicts for the protein-ligand complex.

## Results

### Protein structure modelling and validation

In the modelled protein structure, 91% of the residues are in the most favored region of the Ramachandran plot, 8% of the residues are in the additional allowed region and the remaining 1% of the residues are in the generously allowed region. There are 12 glycine and 27 proline residues in the modelled protein (Fig 1).

**Fig 1:**
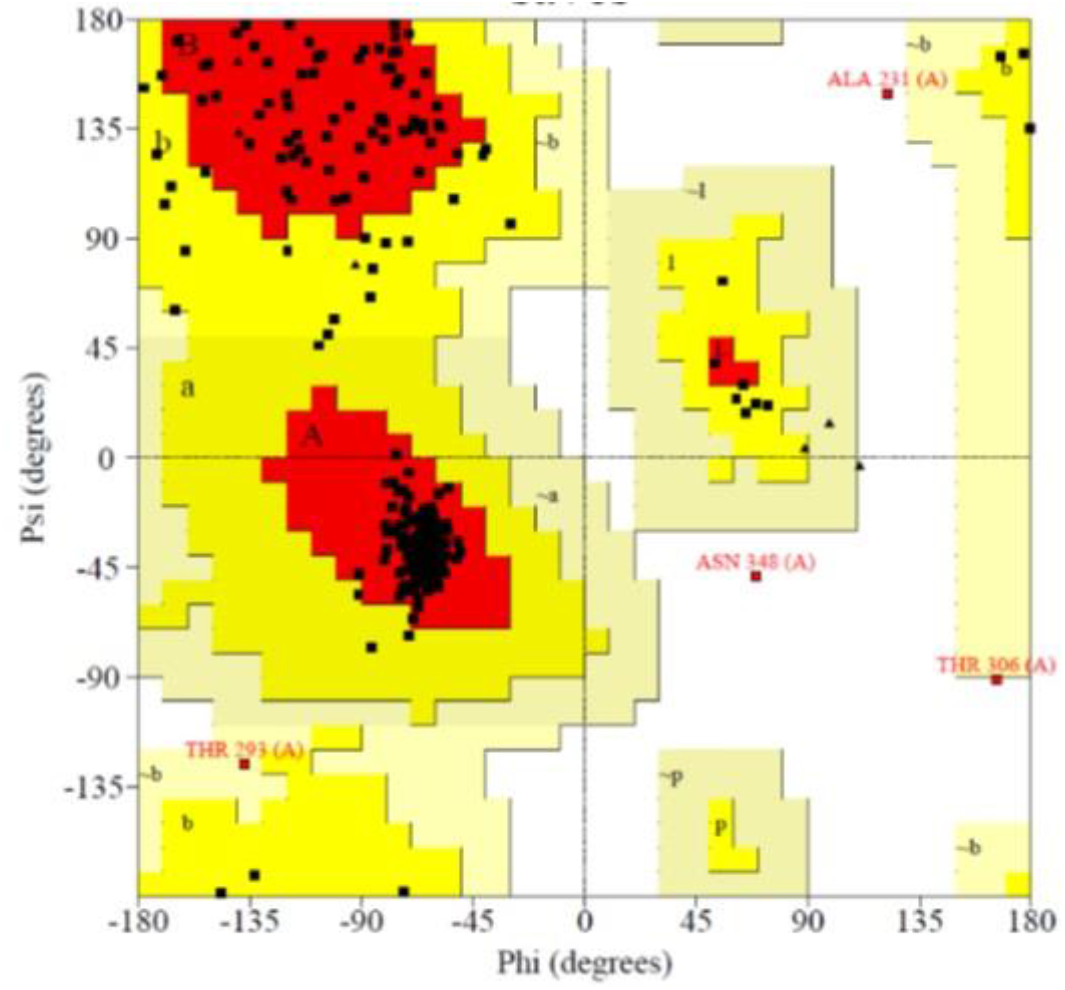
Ramachandran plot for modelled D2 receptor protein. Glycine residues are represented by triangles and proline is represented as ‘p’.

### In-silico molecular docking study for virtual screening of compounds

A molecular docking study is performed to screen and identify potential compounds that show significant binding affinity with the protein of interest and also to identify the nature of the interaction of the compounds with the protein in a protein-ligand docked complex. In our study, we identified 39 phytoconstituents from the IMPPAT database, and these molecules were docked against our modelled D2 receptor protein. In our study, we performed a blind docking approach to identify novel binding sites and this, in turn, enable*s* the ligands to adopt new conformations during the molecular docking study and this flexibility is important to studying various conformational changes of the ligands during docking. Among the 39 phytoconstituents, only 11 molecules with binding energy greater than or equal to -9 kcal/mol are selected for further studies (Table 1).

**Table 1:**
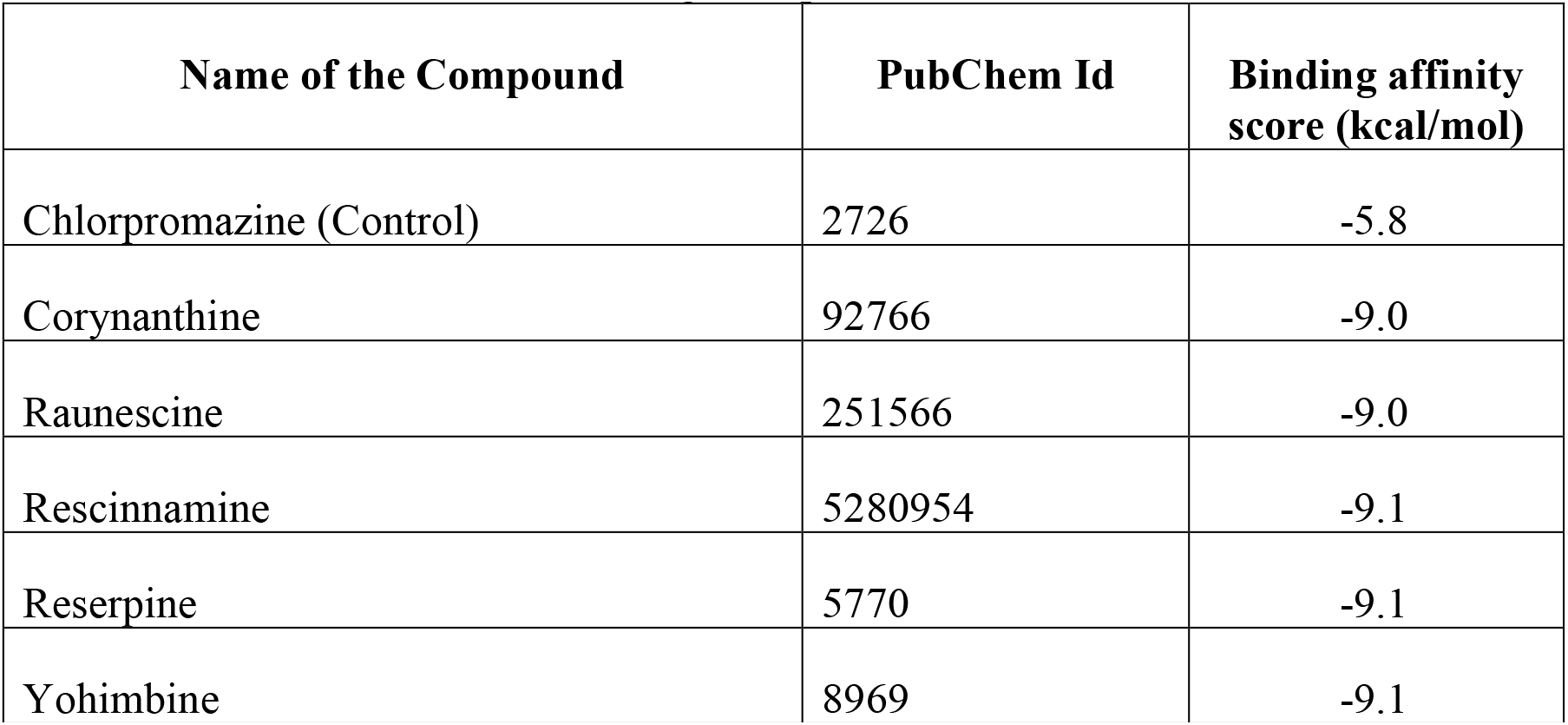
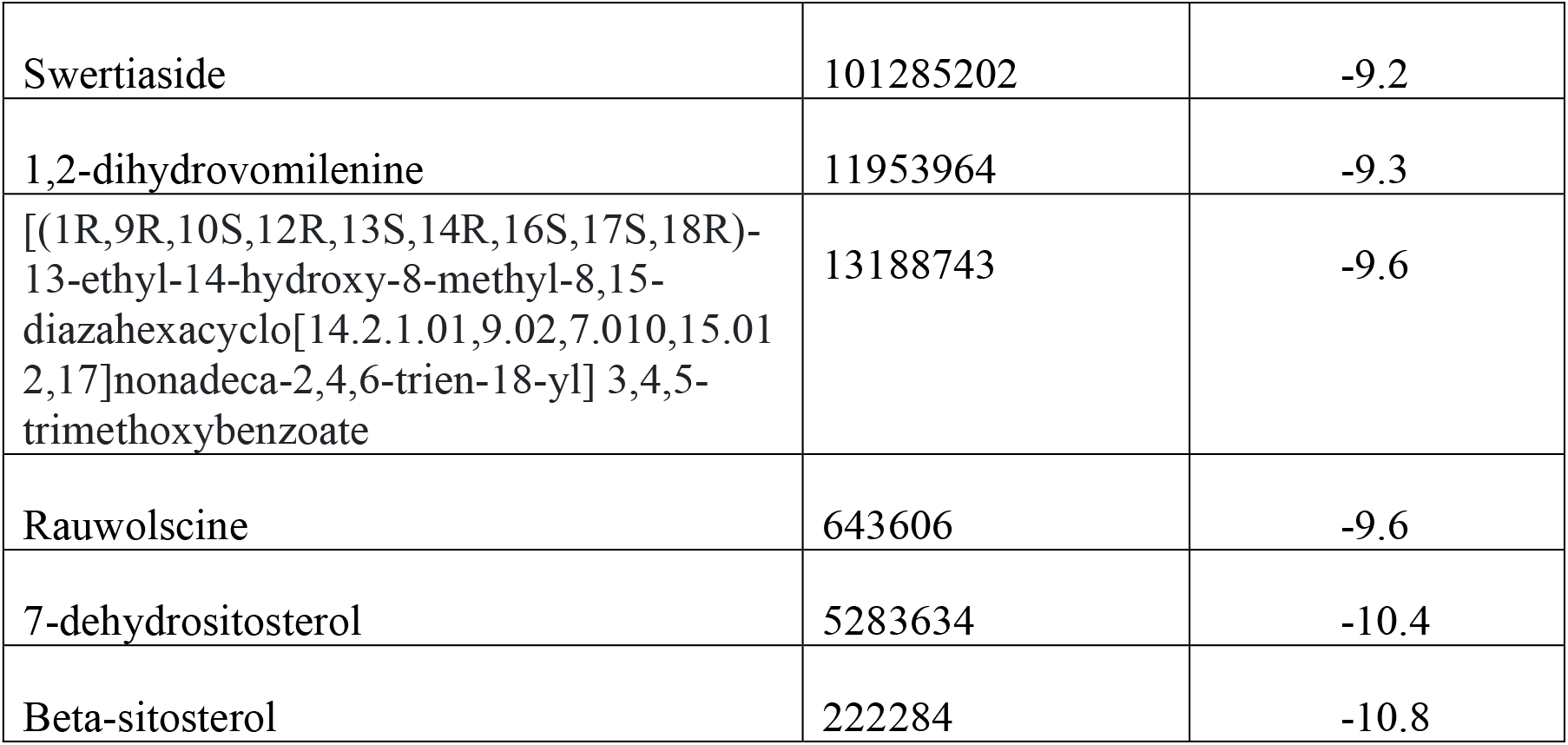
Binding affinity of 11 compounds that passed the ADME filters together with the control drug chlorpromazine.

### ADMET analysis of screened compounds

ADMET analysis refers to the estimation of absorption, distribution, metabolism, and excretion of a particular drug that has been administered to the body. ADMET analysis provides an opportunity to understand the basic pharmacokinetics of a drug under study. For studying the ADMET properties of the identified compounds, we have taken six preset filters as the basis for this analysis and they are the size of the molecule, the solubility of the molecule, the polarity of the molecules, the lipophilicity of the molecule, flexibility and overall saturation of the compound under analysis. In this study we are trying to identify compounds that can enter the brain, therefore for their drug likeliness, Lipinski’s rule of five and blood-brain barrier filters were used and out of 11 compounds, 4 compounds namely 1,2-Dihydrovomilenine (-9.3 kcal/mol), Corynanthine (-9.0 kcal/mol), Rauwolscine (-9.6 kcal/mol) and Yohimbine (-9.1 kcal/mol) passed all the filters discussed above (Fig 2). The toxicity of these four organic molecules was checked to inquire whether these compounds have any tumourigenic, mutagenic effect, and toxic effects of these compounds on our body. The 4 above-mentioned screened compounds passed all the above-set filters without any violations (Fig 2). The side effects of yohimbine are well documented, and. Corynanthine and Rauwolscine are diastereomers of yohimbine and thus toxicity factors have to be tested before these can be considered safe for further study thus these compounds are not considered further. 1,2-Dihydrovomilenine is the only identified compound that does not have any known toxicity risks and has drug-like properties thus 1,2-Dihydrovomilenine docked with D2 receptor protein is considered for further study.

**Fig 2:**
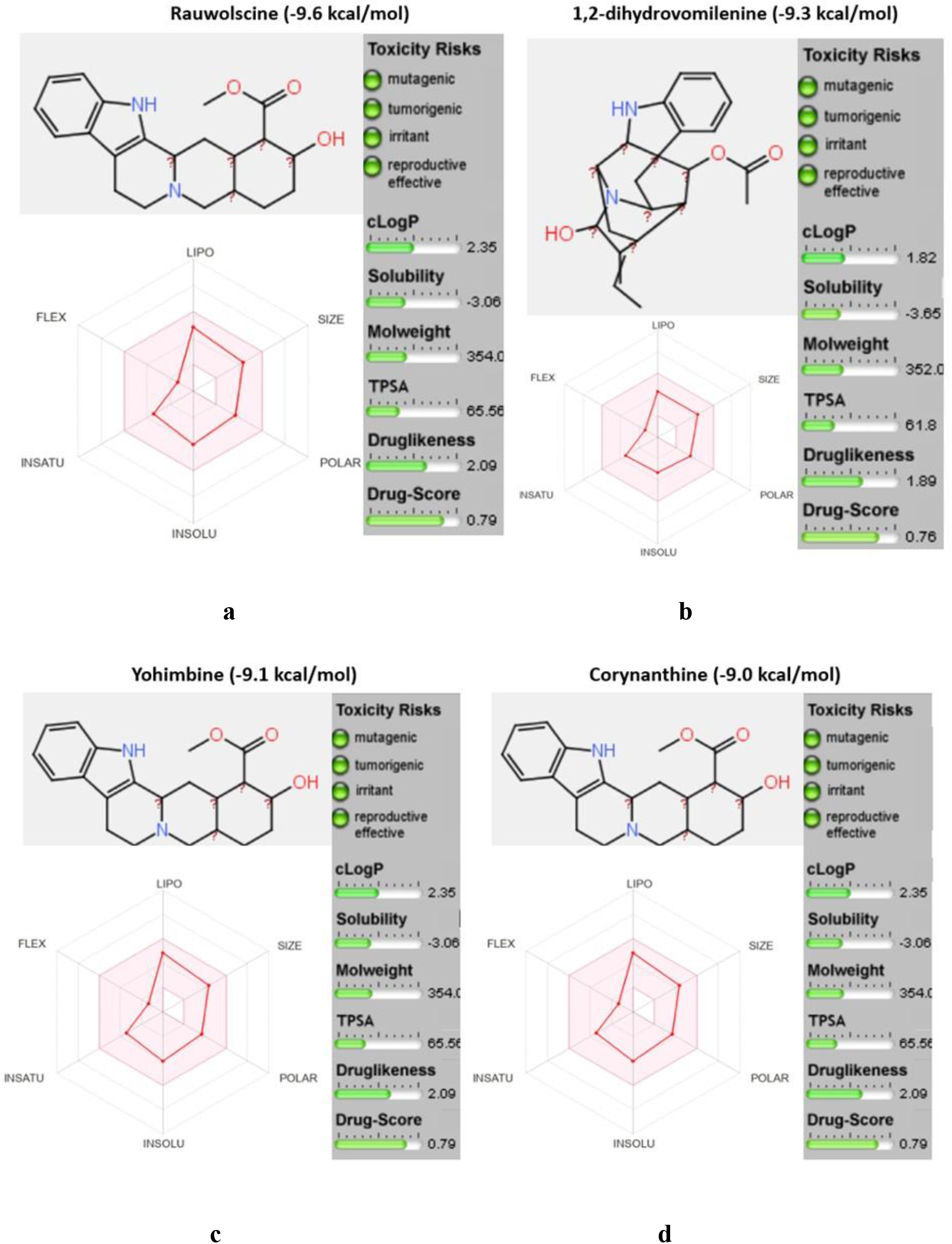
Bioavailability radar together with toxicity profile of Rauwolscine (a), 1,2-dihydrovomilenine (b), Yohimbine (c), and Corynanthine (d). The pink-shaded region of the radar is the estimated physicochemical space for oral bioavailability. The exclamation marks represent carbon with unknown chirality. The safe toxicity profiles of the 4 screened compounds are shown as green tabs in OSIRIS software.

### Visualization of docked complex of D2 receptor with 1,2-dihydrovomelinine

The docked complex of 1,2-dihydrovomelinine with the D2 receptor protein is visualized for studying the interactions of the ligand with the amino acid residues of the protein. The interactions of 1,2-dihydrovomelenine with our protein of interest show that VAL91, LEU94, TRP100, and PHE110 form alkyl and pi-alkyl interaction, TYR408 forms π-π stacked interaction and hydrogen bond with the SER409 residue of the protein (Fig 3).

**Fig 3:**
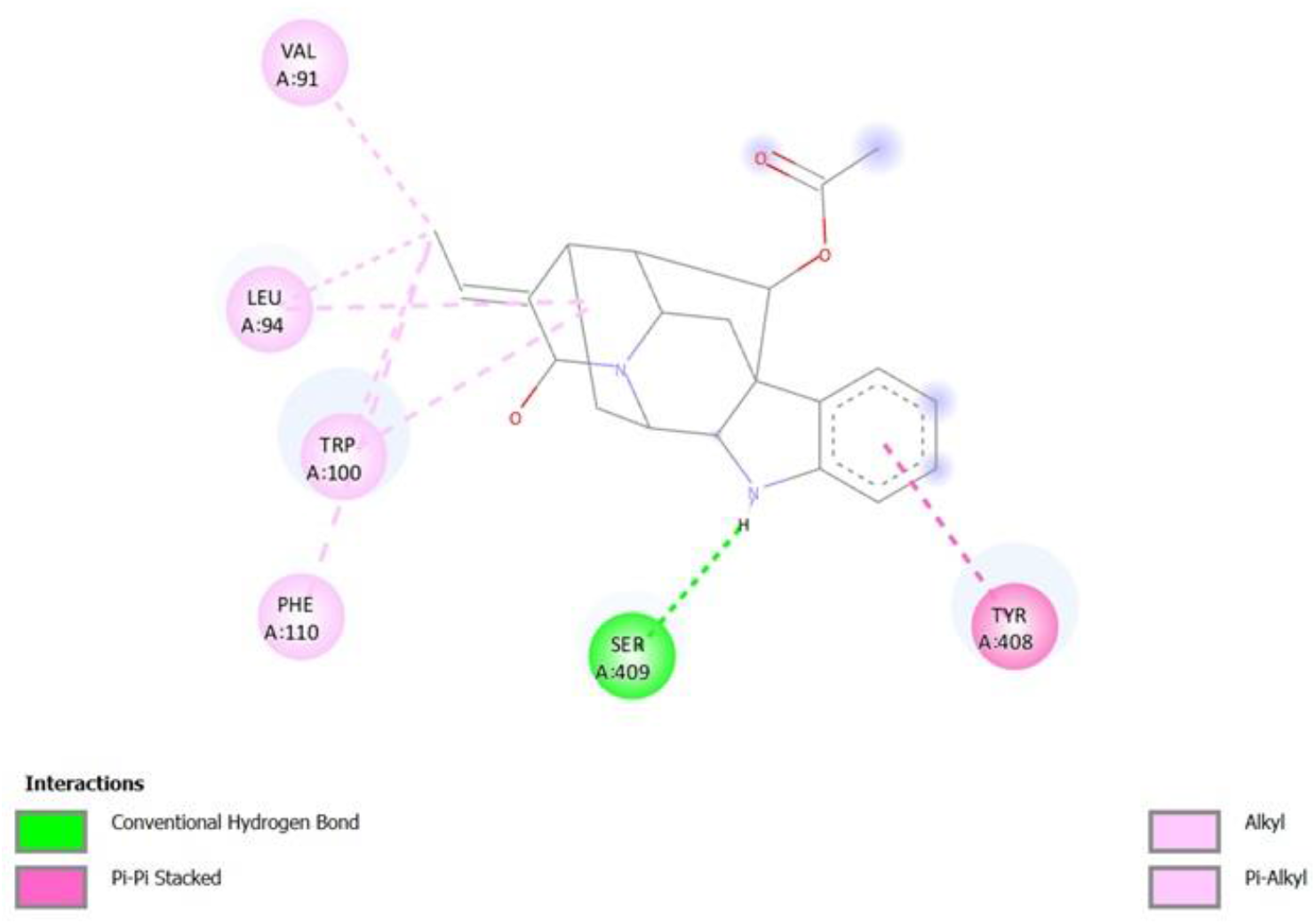
Interaction of 1,2-Dihydrovomilinine with D2 receptor

### Molecular dynamics simulation study of 1,2-dihydrovomelininedocked to D2 receptor

The normal mode analysis of the docked complex showed the deformability of the complex representing the flexibility of the protein from its original conformation. Hinges in protein Fig 4a represent the change in orientation of the protein that shows the interaction of the D2 receptor protein with the ligand. Fig 4b represents the beta factor the complex that represents the displacement of atoms around the state of equilibrium and thus it is equivalent to the root mean square value (RMS) of the protein.

**Fig 4:**
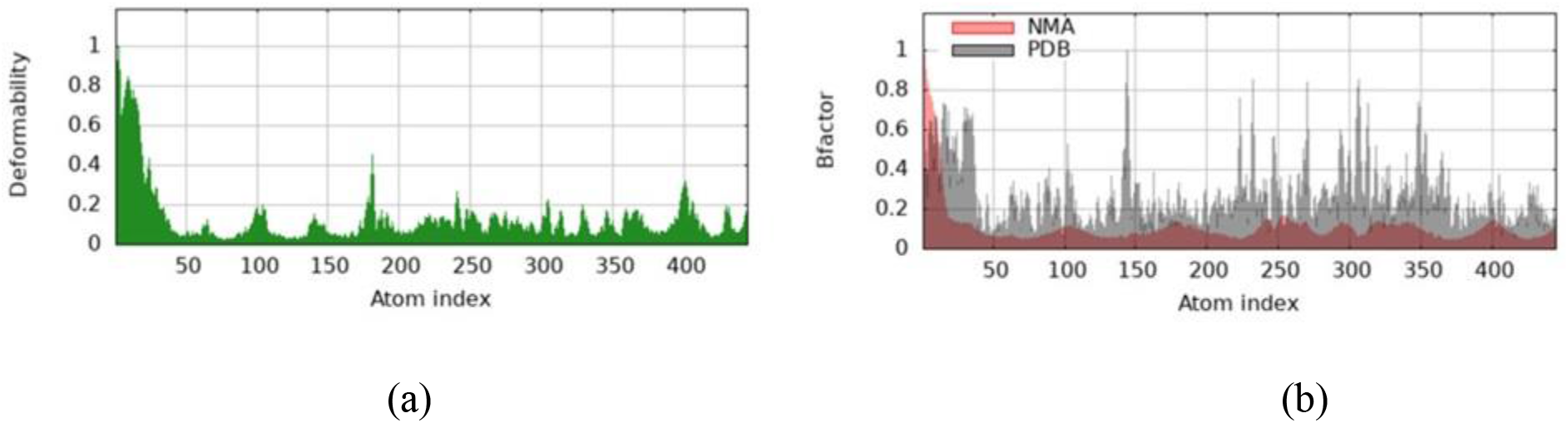
Deformability and beta factor of the docked complex.

## Discussion

D2 dopaminergic receptors are highly expressed in the hippocampus, amygdala, caudate, putamen, substantia nigra, and many other areas of the human brain^23^. D2 receptors serve as a promising target protein for effectively controlling psychotic behaviors and more effectively the positive symptoms that are the hallmarks of Scz, the worst of all kinds of psychotic disorders^24^. In the present study, a molecular docking approach was applied for the virtual screening of bioactive compounds from *Rauwolfia serpentina* from the root parts of the plant. ADMET analysis further streamlined the study leading to the identification of 1,2-Dihydrovomilenine as a potential lead compound (Fig 5). Alkaloids from *R. serpentina* have been studied like ajmaline but the alkaloid identified in this study is a relatively new one and thus it has potential for in-vivo and in-vitro trials^25^. As stated earlier, the D2 dopaminergic receptor has been one of the main targets to control positive symptoms such as hallucinations, and psychosis. Food and Drug Administration (FDA), USA, approved D2 receptor antagonists are divided mainly into first and second-generation antipsychotics. Almost all the first generation antipsychotics such as chlorpromazine, fluphenazine, and haloperidol have some extrapyramidal problems such as dystonia, Parkinsonism and apart from these patients undergoing treatment with these medications can also show metabolic and physiological errors such as weight gain, prolonged QT intervals, low blood pressure^26,27,28^. Second-generation antipsychotics such as risperidone, olanzapine, quetiapine, and ziprasidone shows extrapyramidal side effects together with other side effects such as orthostatic hypotension, increased cholesterol levels, significant weight loss increased cardiac arrythmia, prolonged QT intervals^29,30,31,32^. Because of these above-stated drawbacks, need for an less side effects containing drug is the need of the hour which is the aim of this study.

**Fig 5:**
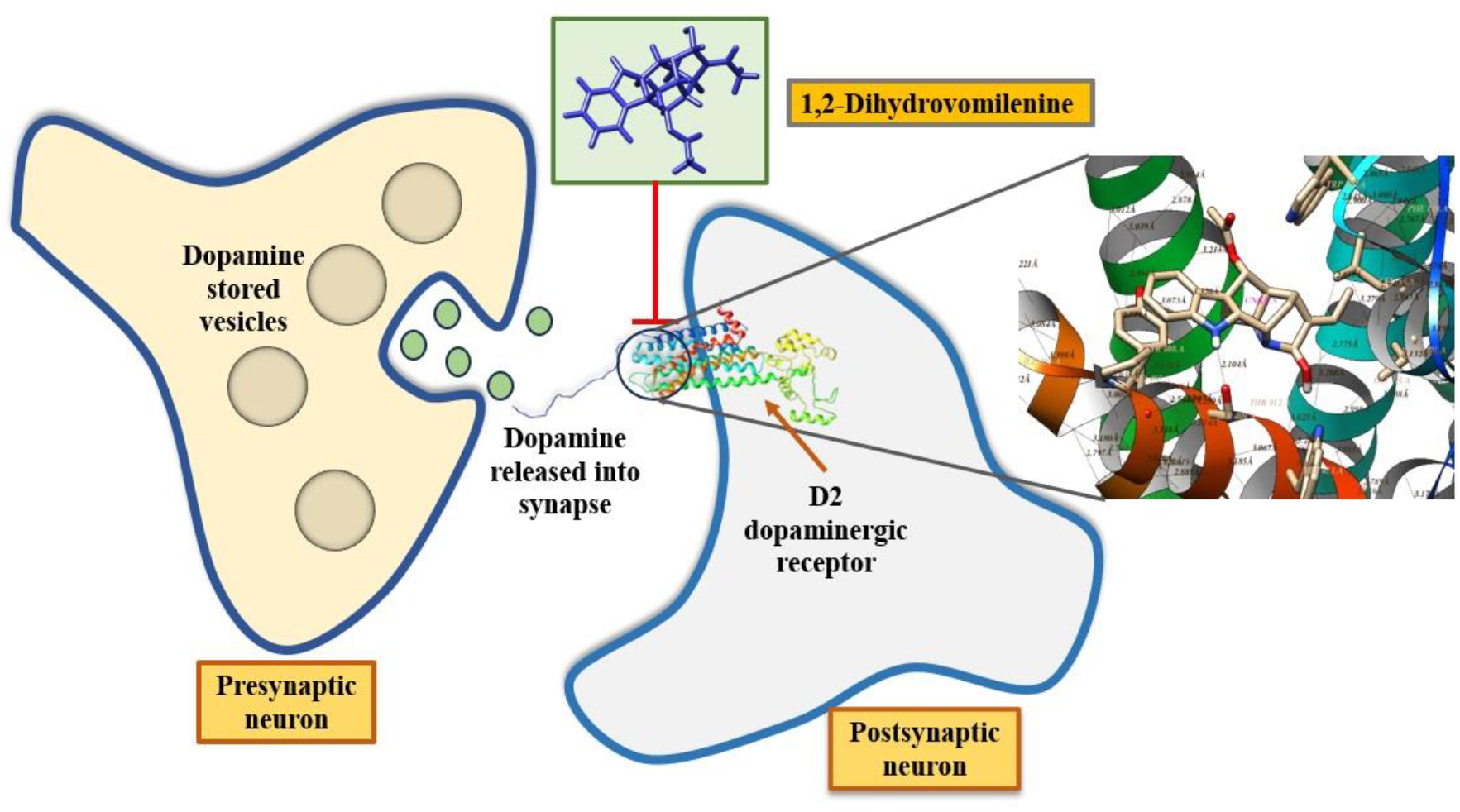
Schematic representation of inhibition of D2 dopaminergic receptor by 1,2-Dihydrovomilenine

All four hits passed the ADME and toxicity filters but out of these four molecules, 1,2-Dihydrovomilenine was chosen as the other three have established cell cytoxicity that has been studied. Out of these three yohimbine has very well documented toxicity factors. In one of the studies, yohimbine was used for physical enhancement but the study concluded that the use of yohimbine did not increase the performance of the bodybuilders also it concluded that in low doses this compound is well tolerated but in case of higher dose toxicity is a factor that should be taken into account^33^. In another study, severe neurotoxicity was detected in a bodybuilder after yohimbine ingestion^34^. Rauwolscine (which is also known as α-yohimbine) and corynanthine are diastereomers of yohimbine and thus their toxicity should be studied in cell lines as well as in the mice models^35^. 1,2-Dihydrovomilenine has 2 hydrogen bond donors and 5 hydrogen bond acceptors according to PubChem and these are good features for becoming a potential drug candidate in the future.

## Conclusion

In this present study, virtual screening of 39 phytochemicals against the D2 dopaminergic receptor has been studied to detect the binding affinity of the phytochemicals for the target protein. According to our study, the 1,2-dihydrovomilenine showed promising results for the target protein and this molecule can be studied further using in-vitro and in-vivo interventions so that it can become a potential drug with high affinity for the D2 receptor with minimum side effects.

## Supporting information

https://drive.google.com/drive/folders/11Db-QXHUtJX53tO2hWG_dXdeq2OXKn3Y?usp=drive_link

